# The alternative sigma factor SigN of *Bacillus subtilis* is intrinsically toxic

**DOI:** 10.1101/2023.03.22.533896

**Authors:** Aisha T. Burton, Debora Kálalová, Elizabeth V. Snider, Andrew M. Burrage, Libor Krásný, Daniel B. Kearns

## Abstract

Sigma factors bind and direct the RNA polymerase core to specific promoter sequences and alternative sigma factors direct transcription of different regulons of genes. Here, we study the pBS32 plasmid-encoded sigma factor SigN of *Bacillus subtilis* to determine how it contributes to DNA damage-induced cell death. We find that SigN causes cell death when expressed at high level and does so in the absence of its regulon suggesting it is intrinsically toxic. One way toxicity was relieved was by curing the pBS32 plasmid, which eliminated a positive feedback loop that lead to SigN hyper-accumulation. Another way toxicity was relieved was through mutating the chromosomally-encoded transcriptional repressor protein AbrB and derepressing a potent antisense transcript that antagonized SigN expression. We note that SigN exhibits a relatively high affinity for the RNA polymerase core, efficiently competing with the vegetative sigma factor SigA, suggesting that toxicity was due to the competitive inhibition of one or more essential transcripts. Why *B. subtilis* encodes a potentially toxic sigma factor is unclear but SigN may be related to phage-like genes also encoded on pBS32.

**SIGNIFICANCE:** Alternative sigma factors activate entire regulons of genes to improve viability in response to environmental stimuli. The pBS32 plasmid-encoded SigN of *Bacillus subtilis* is activated by the DNA damage response and leads to cellular demise. Here we find that SigN impairs viability by hyper-accumulating and outcompeting the vegetative sigma factor for the RNA polymerase core. Why *B. subtilis* retains a plasmid with a deleterious alternative sigma factor is unknown.

## INTRODUCTION

*Bacillus subtilis* is a Gram-positive, model organism due to being domesticated for easy culturing and natural competence which provides facile genetic manipulation (1,2,56). The ancestral strain of *Bacillus subtilis* NCIB3610 is less tractable and carries a large 84 kb plasmid called pBS32 that was lost during the domestication of commonly-used laboratory strain derivatives (3-5). The function of many pBS32-encoded products is unknown, but some have been shown to alter the physiology of the host, including an inhibitor of natural transformation (ComI) (6), an inhibitor of biofilm formation (RapP) (7), and an RNase (RnhP) that contributes to chromosome stability (8). Moreover, nearly half of the episome encodes a large contiguous set of genes resembling a prophage (6), and treatment of cells with the DNA damaging agent mitomycin C (MMC) causes rapid pBS32-dependent cell death (9,10). To date however, no pBS32-derived phage-like particles have been observed after MMC treatment and large deletions of the putative prophage genome do not abolish MMC-dependent death (9). Precisely how and why the plasmid kills the host is unknown.

MMC-induced pBS32-dependent cell death requires the alternative sigma factor SigN encoded on pBS32 (9,10). Expression of SigN is complex and at least three different promoters drive expression of the *sigN* gene. The first promoter (*P_sigN1_*) is repressed by LexA and derepressed by the DNA damage response (10-12). The second promoter is constitutive (*P_sigN2_*) and the third promoter (P*_sigN3_*) is SigN-dependent. Thus when induced by MMC, SigN activates itself and other members of the SigN regulon encoded exclusively on pBS32 (10). Not only is SigN necessary for MMC-induced pBS32-mediated cell death, artificial IPTG-induction of SigN alone is sufficient to cause death when pBS32 is present (9). One way SigN might cause death is if it induced lytic conversion of the putative prophage, but the prophage was not necessary for MMC-induced death and SigN did not appear to activate the prophage structural genes (9,10). Instead, it was presumed that one or more of the genes expressed from the over 20 SigN-dependent promoters on the plasmid was responsible for toxicity. The identity of the toxic gene or genes under SigN control and the mechanism of SigN-dependent cell death was unknown.

Here we explore the mechanism of SigN-mediated cell death by selecting suppressor mutations that restored growth in the presence of artificial SigN induction. Through the analysis of the suppressors mutants in SigN, we found that artificial induction of wild type SigN became toxic even in the absence of pBS32 when expressed at levels higher than that afforded by our standard induction system. Other suppressors spontaneously cured pBS32 and still others mutated AbrB thereby derepressing an anti-sense *sigN* transcript that antagonized SigN accumulation. In sum, we show that at least one toxic product under SigN control is SigN itself. Thus, when the *sigN* transcript exceeds a threshold determined by the anti-sense transcript abundance, the SigN protein is made and initiates a positive feedback loop. SigN toxicity appears to be due to its accumulation combined with its ability to outcompete SigA for RNA polymerase and impair vegetative gene expression. SigN is the founding member of a new subfamily of the sigma 70 family of sigma factors, but the selective advantage, if any, of SigN activity is unclear.

## RESULTS

### Spontaneous suppressors alleviated SigN-mediated cell death

An IPTG-inducible *sigN* construct using the native *sigN* ribosome binding sequence (nRBS) was toxic when introduced to a pBS32-proficient strain such that transformant colonies appeared to be sick even in the absence of IPTG (9). An IPTG-inducible *sigN* construct using a weakened ribosome binding sequence (wkRBS) however, caused a decrease in optical density (OD_600_) only in the presence of IPTG (9,10), but the OD loss was transient, and growth typically recovered 4-5 hours post induction (**Fig 1A**). We inferred that growth recovery was due to genetic suppression. Cultures in which the OD had recovered were dilution plated for single colonies and three distinct classes of <u>s</u>uppressor of *<u>s</u>igN*-mediated <u>d</u>eath (*ssd*) mutants were separated by their colony morphology. Class I mutants had a “ridged” colony morphology and were numerically dominant, whereas class II and III mutants were rare and had either a “wild type” colony morphology or “hyper-rough/hyper-white” colony morphology respectively (**Fig 1B**; **Table 1**). When representatives of each class were isolated and re-challenged with IPTG, cells grew like the wild type and no indication of OD loss was observed. We conclude that SigN-mediated cell death may be genetically suppressed, and we infer that suppressor mutations likely arose in at least three different genetic loci.

**Figure 1:**
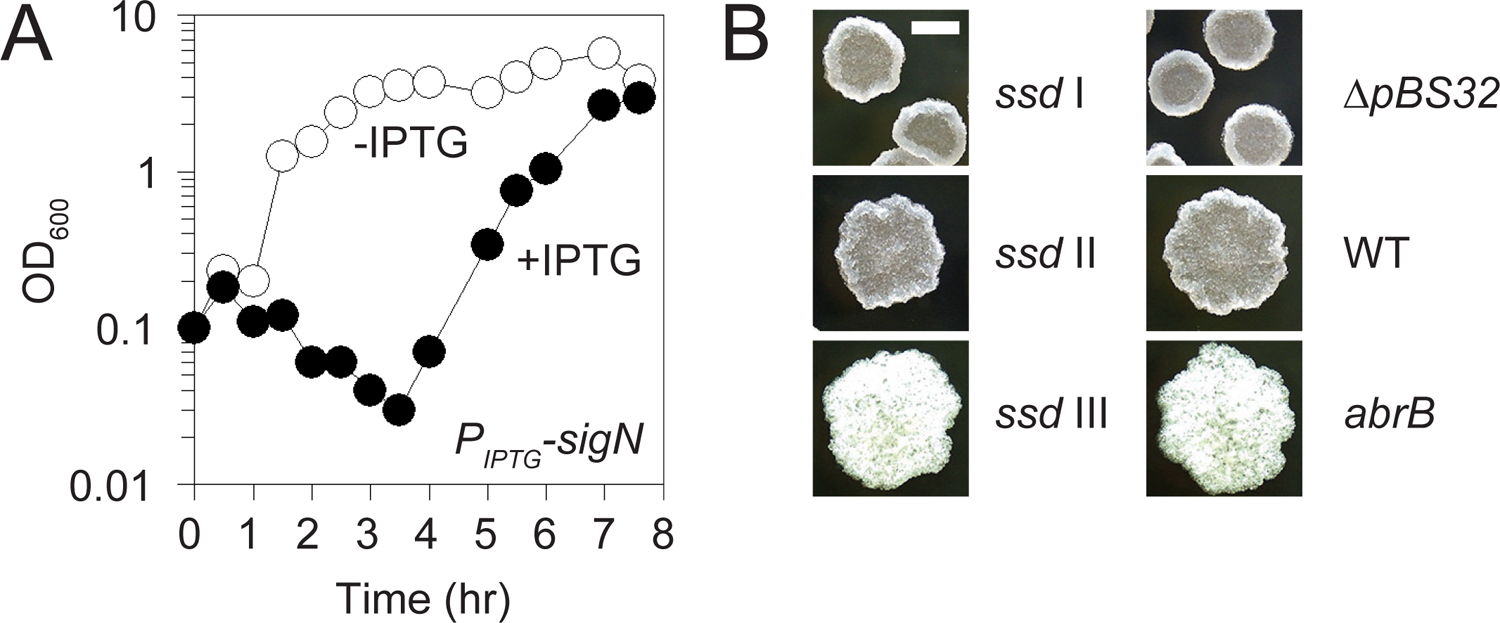
Spontaneous suppressors of SigN-mediated death. A) Optical density (OD_600_) growth curve of a strain containing pBS32 and the ectopic SigN expression construct with a weakened RBS (DK1634) in the absence (open circles) and presence (closed circles) of IPTG. X-axis is the time of spectrophotometry after the addition of IPTG. B) Colony images of *ssd* class I (DK7378), *ssd* class II (DK9591), *ssd* class III (DK6529), *ΔpBS32* (DK451), WT (DK607) and *abrB* (DK5435) mutants. Scale bar is 2 mm.

**Table 1:** Suppressors of SigN-mediated death (*ssd*)

| Mutant | *ssd* Class | Colony<sup>a</sup> | mutation |
| --- | --- | --- | --- |
| DK7378 | I | rough | $\Delta pBS32$ |
| DK7379 | I | rough | $\Delta pBS32$ |
| DK7380 | I | rough | $\Delta pBS32$ |
| DK7381 | I | rough | $\Delta pBS32$ |
| DK7382 | I | rough | $\Delta pBS32$ |
| DK7383 | I | rough | $\Delta pBS32$ |
| DK7384 | I | rough | $\Delta pBS32$ |
| DK7385 | I | rough | $\Delta pBS32$ |
| DK9586 | II | wild type | $P_{hyspank-wkRBS} sigN^{W59*}$ |
| DK9588 | II | wild type | $P_{hyspank-10down-wkRBS} sigN$ |
| DK9589 | II | wild type | $P_{hyspank-wkRBS} sigN^{I198T}$ |
| DK9590 | II | wild type | $P_{hyspank-wkRBS} sigN^{E55G}$ |
| DK9591 | II | wild type | $P_{hyspank-wkRBS} sigN^{K83R}$ |
| DK9592 | II | wild type | $P_{hyspank-wkRBS} sigN^{Q35*}$ |
| DK9282 | IIb | viable | $P_{hyspank-nRBS} sigN^{R22G}$ |
| DK9283 | IIb | viable | $P_{hyspank-nRBS} sigN^{L136P}$ |
| DK9345 | IIb | viable | $P_{hyspank-nRBS} sigN^{D156G}$ |
| DK9342 | IIb | viable | $P_{hyspank-nRBS} sigN^{N160S}$ |
| DK9325 | IIb | viable | $P_{hyspank-nRBS} sigN^{Y165V}$ |
| DK6527 | III | hyper-rough | $abrB^{RBS(AGGAGA>CGGAGA)}$ |
| DK6529 | III | hyper-rough | $abrB^{R15C}$ |
| DK6531 | III | hyper-rough | $abrB^{FS@20bp}$ |
| DK6532 | III | hyper-rough | $abrB^{FS@127bp}$ |

<sup>a</sup>“hyper-rough” colony morphology was distinguishable because it appeared more rugose than the “rough” architecture, was bright white in appearance, and the entire colony had the tendency to leave the agar when prodded with a toothpick.

### Class I *ssd* mutants were cured of pBS32

SigN-mediated cell death was previously shown to require the presence of the pBS32 plasmid (9). Thus, one way in which *ssd* mutations might abolish SigN-mediated cell death is by spontaneous curing of pBS32. To determine whether pBS32 was present in any of the *ssd* suppressor strains, polymerase chain reaction (PCR) was used to amplify three different loci on the plasmid, *zpaB, zpcJ,* and *sigN* (using primers that hybridize to the native site and not the ectopic integration construct), and one locus on the chromosome as a control (*fliG*). Whereas the class II and class III *ssd* mutants amplified all plasmid-encoded loci, the class I *ssd* mutants did not (**Fig 2A**). We conclude that class I *ssd* mutants abolished SigN-mediated cell death by eliminating pBS32 from the genome (**Table 1**). We note that each class I *ssd* allele conferred a ridged colony morphology that phenocopied strains that were force-cured for pBS32 (6,7) (**Fig 1B**).

**Figure 2:**
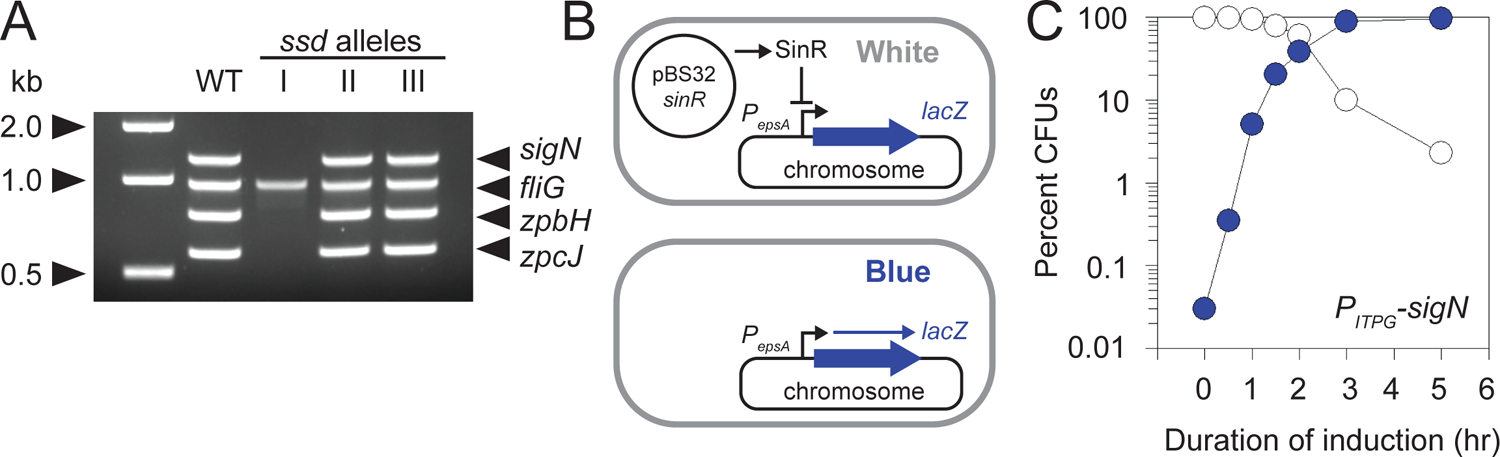
Plasmid curing and mutation of overexpression construct prevent death. A) PCR analysis of representatives of each *ssl* class are shown. Three loci on the plasmid (*sigN*, *zpbH, zpcJ*) and one chromosomal locus (*fliG*) were amplified, the sizes of which are indicated by carets. The NEB 1kb size standard included as a reference in the leftmost lane. Chromosomal DNA from the following strains were used as a template: WT (DK607), *ssd* class I (DK7378), *ssd* class II (DK9591), and *ssd* class III (DK6529). B) Cartoon diagram of the plasmid curing assay. Briefly a chromosomally-encoded β-galactosidase (*lacZ*) reporter is repressed by SinR expressed from the pBS32 plasmid causing colonies to look white when plated on media containing chromogenic substrate X-gal. If the plasmid is cured, SinR is lost and the reporter is de-repressed causing colonies to look blue when plated on media containing X-gal. Thick blue arrow is the *lacZ* gene, thin blue arrow is the *lacZ* transcript, and bent arrow is the SinR-repressed *P_eps_*promoter. Black arrows indicate activation and black T-bars indicate repression. C) Blue/white colony frequency in a pBS32 curing-reporter strain that also encodes a chromosomally encoded IPTG-inducible *sigN* gene with weakened RBS (DK7867). Culture was induced with 1 mM IPTG at time 0, aliquots were taken over time, dilution plated on media containing X-gal, incubated overnight, and counted for the percentage of blue colonies (blue dots) and white colonies (white dots).

Because the class I mutants were cured for the pBS32 plasmid and were the most frequent mutant class, we set out to determine the frequency of plasmid curing in the presence and absence of *sigN* induction. To do so, we constructed a chromosomally-integrated reporter construct in which the *lacZ* gene, encoding the β-galactosidase LacZ, was expressed from the *P_eps_*promoter, the activity of which is de-activated by the biofilm repressor SinR (13,14). Next, we deleted the chromosomal copies of the genes encoding SinR and its antagonist SinI (15), and reintroduced a copy of the *sinR* gene, expressed from its native promoter (16), on the pBS32 plasmid. Thus, when pBS32 expressing SinR was present, *P_eps_-lacZ* activity would be inhibited and colonies would appear white on plates containing the chromogenic substrate, X-gal (**Fig 2B**). If pBS32 were lost however, the gene encoding SinR would also be lost, *P_eps_-lacZ* expression would be de-repressed, and the colony would appear blue.

In the absence of SigN induction, approximately 1 in 3000 colonies of a late exponential phase culture (∼1 OD_600_) was blue when plated on media containing X-Gal indicating that the frequency of spontaneous plasmid curing was roughly 0.03% which is lower than what is noted for the related pLS32 miniplasmid derivative pBET131 (0.5%) (17). Next, IPTG was added to the culture, samples were taken at various durations, and dilution plated on media containing X-gal that lacked inducer. After induction of SigN, the frequency of blue colonies increased exponentially relative to the white colonies (**Fig 2C**). We conclude that cells died after SigN induction but rare cells that had spontaneously cured the pBS32 plasmid prior to induction survived and proliferated. Consistent with the existence of class II and class III suppressors, rare white colonies were observed even after prolonged SigN-induction (**Fig 2C**), suggesting that pBS32 remained in a subpopulation and that there were other mechanisms of SigN-resistance besides plasmid curing.

### Class II *ssd* mutants have mutations in the IPTG-inducible *sigN* construct

Another way in which IPTG-induced SigN-mediated cell death might be abolished is by a loss-of-function mutation in the IPTG-inducible *sigN* construct. To test for *sigN* functionality, the IPTG-inducible *sigN* constructs from the remaining mutants were backcrossed into a wild type background; the resulting strains were induced with IPTG and monitored for growth. Backcrossed IPTG-induced *sigN* alleles of the class II, but not class III, *ssd* mutants failed decrease in OD after induction, indicating that the *sigN* alleles of class II mutants were non-functional. Sequencing of the IPTG-inducible *sigN* gene indicated that one *ssd* allele mutated the −10 element of the IPTG-inducible promoter away from consensus, likely impairing expression (**Table 1**). The remaining five alleles were mutated in the *sigN* open-reading frame including two likely loss-of-function nonsense mutations (SigN^W59stop^ and SigN^Q35stop^) and three missense mutations (SigN^E55G^, SigN^K83R^, and SigN^I198T^) (**Table 1**). We conclude that the class II *ssd* mutants eliminated SigN-mediated cell death by either reducing or abolishing SigN expression or activity.

To determine whether the missense mutations in *sigN* impaired or abolished SigN accumulation, we set out to detect SigN protein in lysates. After 2 hours of IPTG induction, levels of SigN protein were high in the wild type but reduced in the *sigN* mutants (**Fig 3A**). While the three mutant substitutions might have each resulted in unstable proteins, we were concerned that they might also be reduced due to impaired activity. The IPTG-inducible SigN construct was built with a weakened RBS (wkRBS) to restrict activity and substantial amounts of SigN might only be achieved when SigN auto-activates at the *sigN* gene located on pBS32. Thus, high levels of SigN protein may require SigN functionality as non-functional alleles, expressed from the weakened RBS, might fail to stimulate positive feedback at the native site. We conclude that the IPTG-inducible SigN mutant alleles produced low levels of protein but precisely why protein levels were low was uncertain.

**Figure 3.**
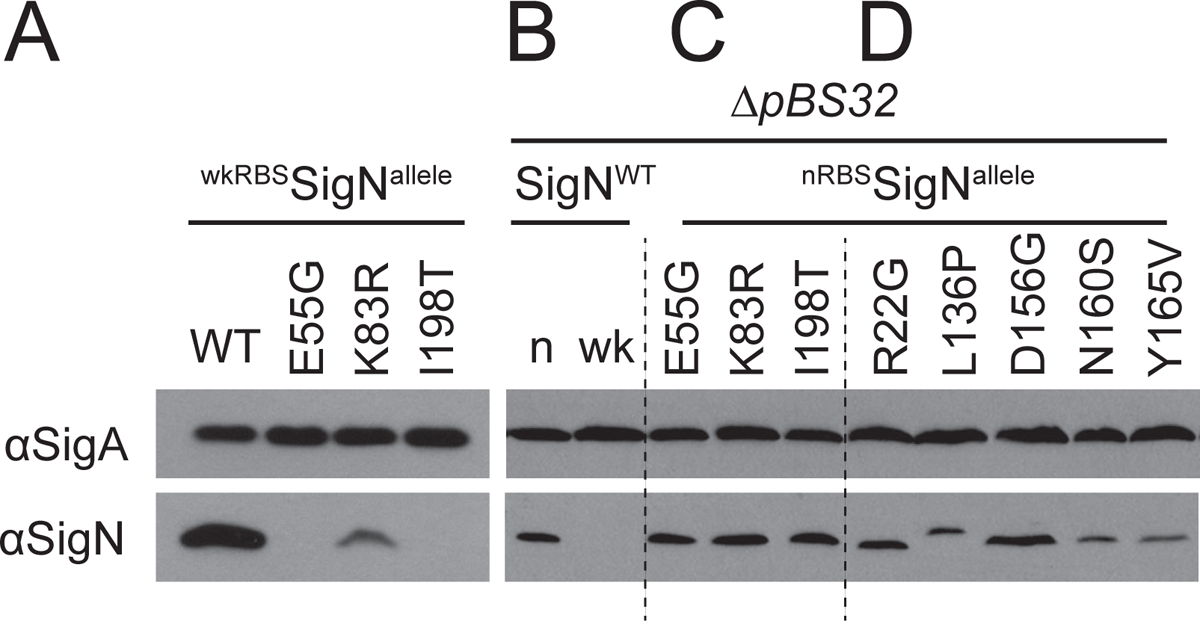
Missense mutants that alter the SigN sequence produce levels of protein like the wild type. Western blot analysis of whole cell lysates detected with anti-SigA as a loading control (top panel) and anti-SigN (bottom panel) is shown. A) Wild type SigN protein (DK1634) or the indicated SigN alleles (DK9590, DK9591, DK9589) were expressed from an IPTG-inducible promoter with a weakened RBS “wk” in a strain containing pBS32. B) Wild type SigN protein was expressed from an IPTG-inducible promoter with either the native SigN RBS “n” (DK9067), or a weakened RBS “wk” in a strain lacking pBS32 (DK9069). C) Each of the indicated SigN amino acid substitution mutants selected to abolish pBS32-dependent cell death were expressed from an IPTG-inducible promoter and native SigN RBS (^nRBS^SigN) in a strain lacking pBS32. The following strains were used to generate this panel: DK9210 (SigN^E55G^), DK9211 (SigN^K83R^), and DK9209 (SigN^I198T^) D) Each of the indicated SigN amino acid substitution mutants selected to abolish pBS32-independent cell death were expressed from an IPTG-inducible promoter and native SigN RBS (^nRBS^SigN) in a strain lacking pBS32. The following strains were used to generate this panel: DK9282 (SigN^R22G^), DK9283 (SigN^L136P^), DK9345 (SigN^D156G^), DK9342 (SigN^N160S^), and DK9325 (SigN^Y165V^) For each lane, cells were induced at 0.5 OD_600_ or less with 1 mM IPTG for 2 hours prior to harvesting cell pellets for lysis. Panels B, C, and D are from the same gel and are separated by dashed lines to facilitate citation in text.

In a second attempt to detect SigN mutant proteins, the wild type and mutant alleles were expressed from an IPTG-inducible promoter using the native SigN RBS (nRBS) sequence and inserted at an ectopic site in a laboratory strain lacking pBS32. In the absence of pBS32, integration of the IPTG-inducible wild type *sigN* allele with nRBS did not appear to impair transformation nor did it appear to reduce growth rate in the absence of IPTG. Moreover, the SigN protein was detected but only when expressed using the nRBS sequence, supporting the notion that the wkRBS construct required positive feedback on pBS32 to produce detectable levels (**Fig 3B**). We noted however that expression of SigN with the nRBS inhibited cell growth even in the strain lacking pBS32 whereas the SigN with the wkRBS did not (**Fig 4A**). We conclude that expression of SigN from its nRBS is sufficient to produce detectable levels of protein in the absence of positive feedback from the pBS32 plasmid. We further conclude that detectable levels of SigN protein resulted in growth inhibition by a pBS32-independent mechanism.

**Figure 4:**
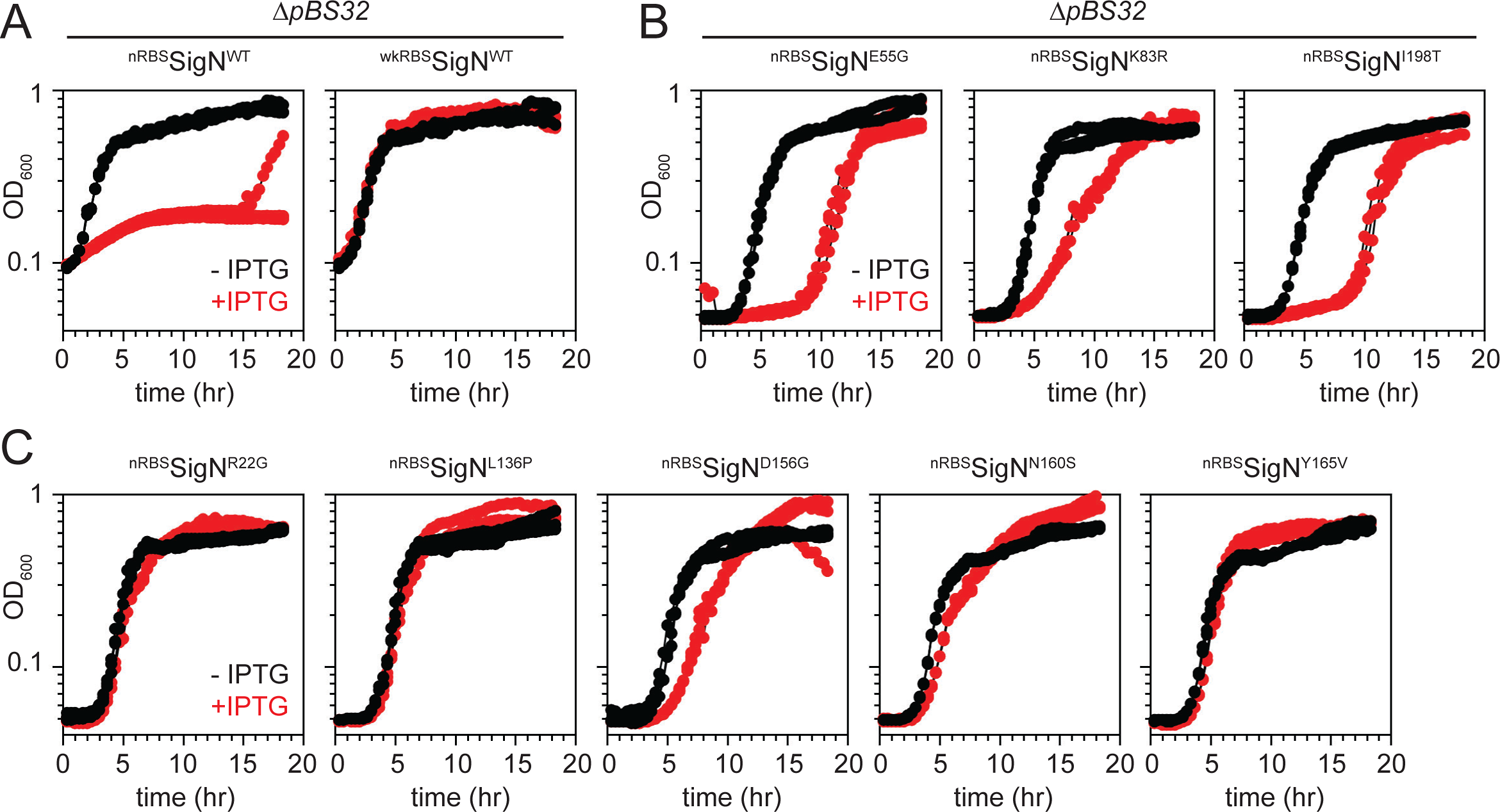
Expression of SigN from its native RBS arrests growth in the absence of pBS32. A) Plate reader growth curves of strains containing an IPTG-inducible SigN with either native RBS (nRBS, DK9067) or weakened RBS (wkRBS, DK9069) grown in the presence (red) and absence (black) of 1 mM IPTG. Both strains lack the pBS32 plasmid. B) Plate reader growth curves of strains containing IPTG-inducible SigN^E55G^ (DK9210), SigN^K83R^ (DK9211), or SigN^I198T^ (DK9209) with native RBS grown in the presence (red) and absence (black) of 1 mM IPTG. C) Plate reader growth curves of strains containing IPTG-inducible SigN^R22G^ (DK9282), SigN^L136P^ (DK9283), SigN^D156G^ (DK9345), SigN^N160S^ (DK9342), or SigN^Y165V^ (DK9325) with native RBS grown in the presence (red) and absence (black) of 1 mM IPTG. Each strain was grown in triplicate and each growth curve is presented separately.

Next, each of the class II *sigN* alleles were artificially expressed with the nRBS in a strain lacking pBS32. Induction of each of the class II missense alleles produced levels of SigN protein comparable to the wild type suggesting that they were each likely defective in SigN function, rather than SigN protein stability (**Fig 3C**). The alleles were not completely defective in SigN activity however as induction of each of mutant allele at least partially inhibited growth in the absence of pBS32 (**Fig 4B**). We conclude the class II mutant alleles expressed from the wkRBS abolished cell death in the presence of pBS32 because they lacked an activity of SigN necessary to initiate positive feedback. We further conclude that the alleles nonetheless retained an activity that caused pBS32-independent growth inhibition when their levels were raised by other means.

To isolate mutant variants of SigN which abolished pBS32-independent growth inhibition, the IPTG-inducible *^nRBS^sigN* construct was grown to high density in the absence of inducer and then plated on media containing IPTG. Rare colonies survived, five were clonally isolated, and sequencing revealed that each strain contained a mutation in the *sigN* open reading frame. Each of the new alleles that abolished pBS32-independent cell death also abolished death when crossed into a background containing pBS32, and were therefore classified as a subset (“b”) of *ssd* class II alleles (**Fig 4C, Table 1**). Finally, the level of SigN protein from each *ssd* class IIb allele was comparable to the wild type in Western blot analysis (**Fig 3C**). We conclude that pBS32-independent death and pBS32-dependent death activities are related as SigN mutants defective in the former also abolished the latter. We infer that SigN likely inhibits growth by a single mechanism and that the pBS32-dependent phenotypes require the plasmid to create positive feedback on SigN expression.

The SigN regulon is restricted to pBS32 (10). As SigN does not appear to stimulate transcription from the chromosome, the mechanism of pBS32-independent cell death is likely unrelated to expression of a chromosomal gene and more likely due to intrinsic inhibition by the SigN protein itself. To determine whether SigN might function as a generalized transcription inhibitor, SigN was added to an *in vitro* transcription assay containing the RNA polymerase core, the vegetative sigma factor SigA, and a DNA fragment containing SigA-dependent (*P_sigN2_*) and SigN-dependent (*P_sigN3_*) promoters from upstream of *sigN* cloned into the transcription vector p770 (Ross 1990). In the absence of SigN, SigA-containing RNA polymerase synthesized the SigA-dependent transcript (**Fig 5A**). As SigN levels rose in the reaction, SigN-dependent transcripts increased, while SigA-dependent transcripts decreased such that a two-fold excess of SigN was sufficient to inhibit SigA transcription by 50%. Increasing the amount of the alternative stress response sigma factor SigB did not reduce the SigA-dependent transcript from the same template as much, suggesting that SigN was a stronger inhibitor and better competitor for the RNAP core (**Fig 5B, Fig S1**) (Rollenhagen 2003, Vohradsky et al., 2021). We conclude that SigN is a potent competitive inhibitor of the vegetative sigma factor and we infer that SigN-mediated growth inhibition may be due to a reduction of essential vegetative transcripts.

**Figure 5.**
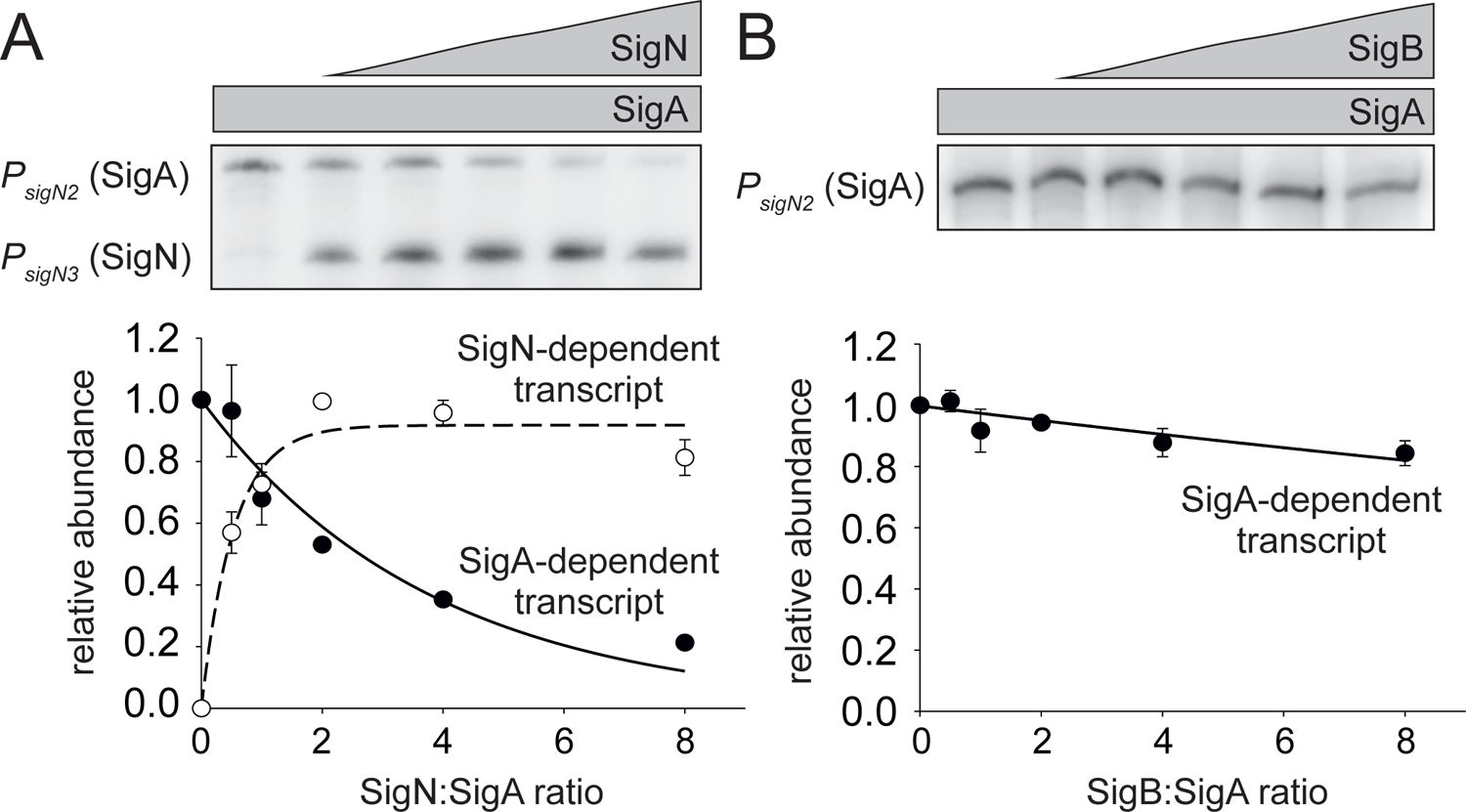
Relative affinities of *B. subtilis* SigA, SigN and SigB for the RNAP core. Multiple-round transcriptions were carried out with increasing ratios of alternative sigma factors to SigA. The DNA template contained two promoters: P2 (*P_sigN2_* SigA-dependent; longer transcript) + P3 (*P_sigN3_* SigN-dependent; shorter transcript). Representative primary data are shown above the graphs. Individual data points are averages from at least three biological replicates, the error bars indicate standard deviations of density scans. (**A**) Competition between SigA and SigN for the RNAP core. SigA (open circles) and SigN transcripts (closed circles) were detected. Transcription from P2 in the absence of SigN was set as 1 (first lane). (**B**) Competition between SigA and SigB for the RNAP core. Transcription from P*_sigN2_* in the absence of SigB was set as 1 (first lane).

### Class III *ssd* alleles have mutations in the transcriptional repressor, AbrB

The class III *ssd* mutants were not mutated for the inducible *sigN* construct nor were they cured of pBS32. We inferred that the class III mutants contained loss-of-function mutations like the class II mutants, because the two classes occurred at approximately the same rare frequency in the spontaneously suppressed population. To find the location of the class III mutations, a strain containing the IPTG-inducible *P_hyspank_-^wkRBS^sigN* construct was mutagenized with a *mariner* transposon carrying a chloramphenicol resistance cassette, and the mutagenized pool was plated on media supplemented with both chloramphenicol and IPTG. Some of the rare colonies that grew had a hyper-rough/hyper-white colony morphology (like the spontaneous class III mutants), two of which were isolated and backcrossed to ensure linkage between the chloramphenicol resistance, colony morphology phenotype, and survival in the presence of IPTG. Finally, the location of the transposon was determined and for each strain, the transposon was inserted within the chromosomally encoded gene, *abrB* (**Fig 1B, Fig 6A**). Sequencing of *abrB* in each of the class III *ssd* mutants indicated the presence of mutations within the gene that altered either the protein sequence or likely reduced translation of the gene product (**Table 1**).

**Figure 6:**
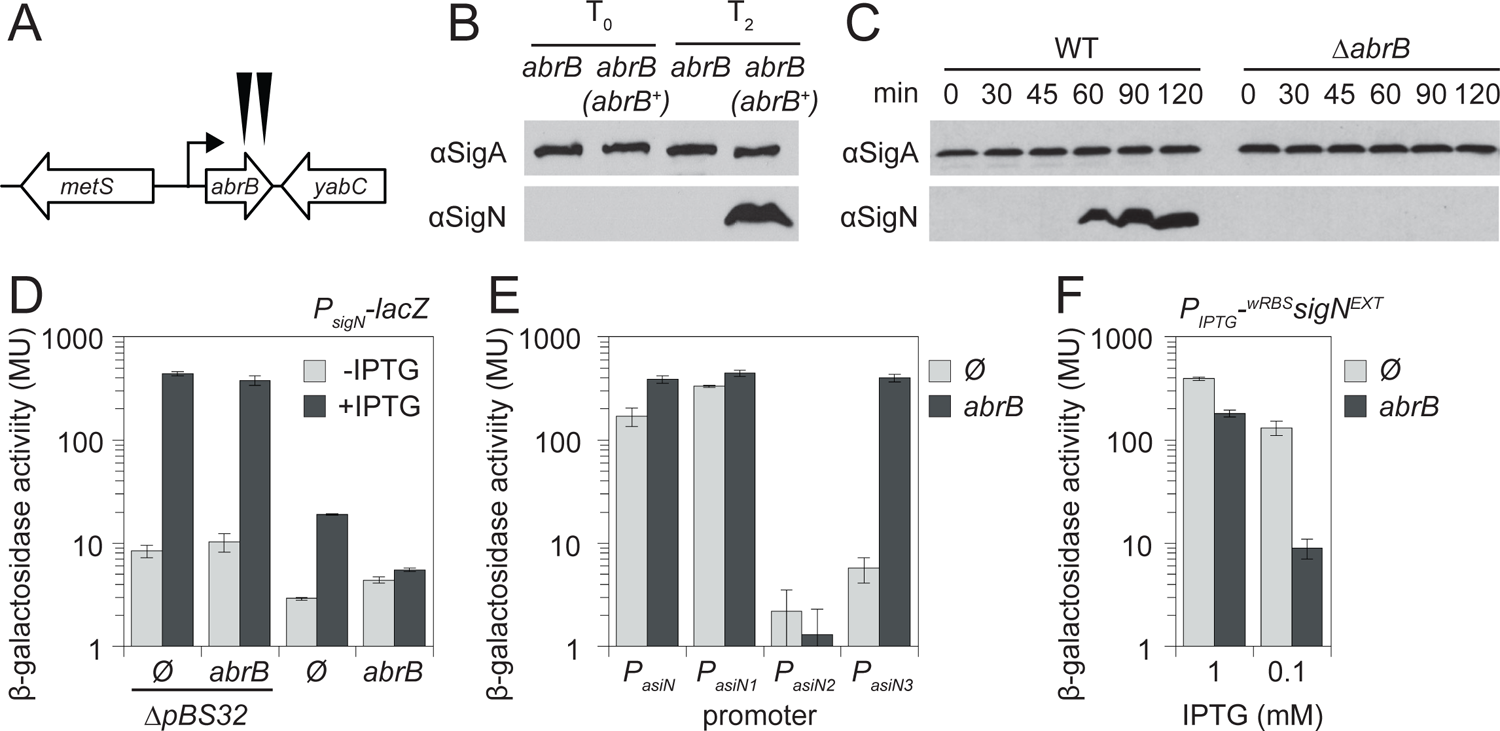
Mutation of AbrB suppresses SigN-mediated cell death. A) Cartoon diagram of the *abrB* genetic neighborhood. Open arrow indicate genes, bent arrow indicates promoter. Black carets indicate the location of transposon insertions in *abrB* (DK3140 and DK3144). B) Western blot of SigN accumulation when ^wkRBS^SigN was induced immediately after IPTG addition (T_0_) and two hours afterwards (T_2_) in an *abrB* mutant (DK6955) and an *abrB* mutant complemented with *abrB* ectopically expressed from its native promoter (*abrB^+^*) (DK7132). The vegetative sigma factor SigA was also included as a control. C) Western blot of SigN accumulation immediately after the indicated times following addition of the DNA damaging agent mitomycin C in otherwise wild type (DK607) and an *abrB* mutant (DK5435). The vegetative sigma factor SigA was also included as a control. D) β-galactosidase assays of strains containing an IPTG-inducible *^wkRBS^sigN* gene and a *P_sigN_-lacZ* reporter grown in the absence (light gray bars) and presence (light gray bars) of 1 mM IPTG for 1 hour. Ø indicates an otherwise wild type background, *ΔpBS32* indicates strains in which the pBS32 plasmid has been cured. The following strains were used to generate this panel: Ø *ΔpBS32* (DB250), *abrB ΔpBS32* (DB251), Ø (DB249), and *abrB* (DB261). Error bars are the standard deviations of three replicates. E) β-galactosidase assays of strains containing a *lacZ* reporter fused to the promoter region indicated on the x-axis. Each strain contained either an *abrB* mutation (dark gray bars) or no additional modification (Ø, light gray bars). The following strains were used to generate this panel: *P_asiN_-lacZ* Ø (DB311) and *abrB* (DB307), *P_asiN1_-lacZ* Ø (DB312) and *abrB* (DB308), *P_asiN2_-lacZ* Ø (DB313) and *abrB* (DB309), *P_asiN3_-lacZ* Ø (DB314) and *abrB* (DB310). Error bars are the standard deviations of three replicates. F) β-galactosidase assays of *ΔpBS32* otherwise wild type (Ø, light gray bars, DB259) or *abrB* mutant (dark gray bars, DB260) strains containing an IPTG-inducible *^wkRBS^sigN^EXT^* gene with extended downstream region and a *P_sigN_-lacZ* reporter grown in the presence of either 1 mM IPTG or 0.1 mM IPTG for 1 hour. Error bars are the standard deviations of three replicates.

The *abrB* gene encodes AbrB, a DNA binding transcription factor (20,21) and a known transcriptional repressor of biofilm development and antimicrobial production (22-24). One way in which AbrB might alleviate SigN-induced cell death is if mutation of *abrB* abolished or reduced SigN expression. Consistent with a SigN expression defect, no SigN was detected in an *abrB* mutant after ^wkRBS^SigN was induced for two hours in the presence of pBS32, but SigN protein was detected when *abrB* was complemented under the control of its native promoter and integrated at an ectopic site in the chromosome (**Fig 6B**). Moreover, SigN protein accumulated in the wild type two hours after induction from the native site by treatment with the DNA damaging agent mitomycin C (MMC), and MMC-dependent accumulation was abolished in the *abrB* mutant (**Fig 6C**). We conclude that mutation of *abrB* prevents SigN-mediated death by preventing SigN accumulation, and does so whether SigN is induced artificially by IPTG or by the DNA damage response at the native site in the chromosome.

One way that mutation of *abrB* could impair accumulation of SigN is if it altered *sigN* transcription. To measure *sigN* transcription, a reporter in which the *P_sigN_* promoter region was fused to the *lacZ* gene encoding β-galactosidase was introduced to a strain with an IPTG-inducible ^wkRBS^*sigN* construct and lacking pBS32. Consistent with previous reports, IPTG-induction of *sigN* induced expression of the *P_sigN_-lacZ* reporter over 50-fold (10), and mutation of *abrB* did not diminish *P_sigN_* induction (**Fig 6D**). We conclude that AbrB does not directly act upon the *P_sigN_* promoter nor does it inhibit SigN activity. To determine whether AbrB might inhibit SigN activity indirectly through a pBS32-encoded product, the reporter expression experiments were conducted in a pBS32-containing strain. Consistent with an indirect effect, induction of the *P_sigN_*promoter was reduced when pBS32 was present, and mutation of AbrB abolished induction entirely (**Fig 6D**). We conclude that AbrB indirectly activates SigN and does so in a manner that depends on a pBS32-derived product.

One possible candidate for a regulator of SigN is a putative antisense RNA initiated downstream of, and reverse transcribing through, the *sigN* gene (25). To determine whether a promoter resides downstream of, and oriented towards, the *sigN* gene, the putative anti-*sigN* promoter region (*P_asiN_*) was cloned upstream of the *lacZ* gene and inserted at an ectopic site in the chromosome. Consistent with the presence of one or more functional promoters, β-galactosidase activity was detected from the *P_asiN_* region and expression from *P_asiN_* increased when AbrB was mutated (**Fig 6E**). REND-seq predicted three different possible start sites that could correspond to different promoter elements and the *P_asiN_*region was divided into three separate fragments, *P_asiN1_*, *P_asiN2_*, and *P_asiN3_* each cloned separately upstream of the *lacZ* gene (**Fig 7**). The *P_asiN1_*promoter expressed at a high level and appeared constitutive, while the *P_asiN2_* putative promoter gave no activity either in wild type or in an *abrB* mutant (**Fig 6E**). Consistent with repression however, the *P_asiN3_* promoter produced very low transcript in wild type but expression increased 100-fold when *abrB* was mutated (**Fig 6E**). We conclude that at least two promoters drive anti-sense *sigN* transcription, one of which is constitutive and one of which is tightly repressed by AbrB.

**Figure 7.**
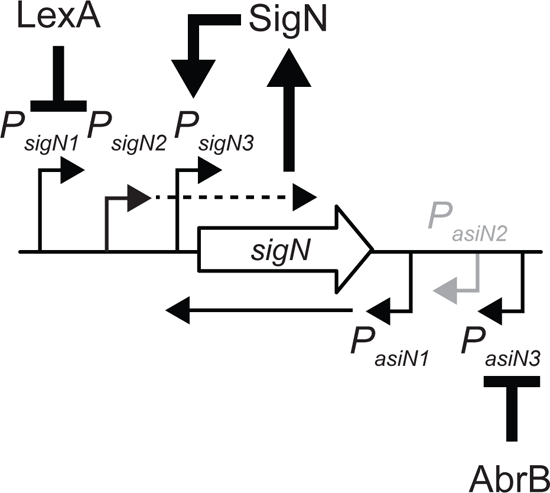
Model of SigN transcriptional regulation. Cartoon diagram of the *sigN* genetic neighborhood. Open arrow indicate genes, bent arrow indicates promoter. Heavy arrows indicate activation. Heavy T-bars indicate repression.

Each of the ectopic, inducible *sigN* constructs used thus far were built excluding the region downstream of *sigN* that contains the anti-sense promoters. To determine the biological relevance of the antisense transcript, the *sigN* gene was cloned with a wkRBS and the extended downstream region under the control of an IPTG inducible promoter (*P_hypsank_-^wkRBS^sigN^EXT^*). Next the expression construct was inserted at an ectopic site in strains containing the *P_sigN_-lacZ* reporter but lacking pBS32 (to eliminate the contribution of natively produced antisense transcript). Expression from *P_sigN_* decreased slightly in the *abrB* mutant when SigN was induced with 1 mM IPTG, but expression decreased dramatically when the amount of inducer was reduced 10-fold (**Fig 6F**). We conclude that the AbrB effect was at least partially linked to the region downstream of *sigN*. We further conclude that de-repression of the *P_asiN3_* promoter and production of the antisense *sigN* transcript was able to restrict SigN to levels insufficient for positive feedback thereby avoiding sequestration of the pool of RNA polymerase core.

## DISSCUSSION

SigN appears to be the founding member of a new sub-family of sigma 70 sigma factors, and SigN autoactivates its own expression on pBS32 (10). Here we set out to determine which gene or genes under SigN control inhibited growth in a pBS32-dependent manner, and discovered that at least one of those genes was SigN itself as SigN was intrinsically toxic in the absence of the rest of the regulon. Toxicity is likely due to the ability of SigN to outcompete the vegetative SigA for RNA polymerase core leading, and competition may be further exacerbated by DNA relaxation after DNA damage (26,27). Two different classes of loss-of-function mutation in SigN were found that differentially reduced toxicity. One class relieved pBS32-depedent inhibition and we speculate that these altered residues that are involved in promoter recognition. The other class relieved pBS32-independent inhibition and we speculate that these altered residues involved in interaction with the RNA polymerase core. *In vivo*, the two activities direct positive feedback at *sigN*, causing SigN to accumulate and increase competition with SigA. Thus, SigN likely inhibits growth by sequestering the RNA polymerase core and reducing expression of one or more essential genes to sub-viable levels. If true, one might expect SigN auto-activation to be tightly regulated and indeed, its levels are restricted at both transcriptional and post-transcriptional levels.

One way to control SigN levels is transcriptional by its complex native promoter region including a basal constitutive promoter (*P_sigN2_*), a promoter de-repressed by the DNA-damage response (*P_sigN1_*), and a promoter that is SigN-dependent (*P_sigN3_*) (**Fig 7**). Normally, the constitutive promoter is insufficient to initiate positive feedback and requires the DNA-damage response to push SigN expression above the feedback threshold (10). The threshold however could also be exceeded artificially, and the original report of SigN suggested that even expression from an uninduced IPTG-inducible promoter was sufficient to cause morbidity (9). Accordingly, the SigN RBS was weakened to make toxicity dependent on both IPTG and pBS32, and here we show that perhaps the only reason pBS32 was needed was to provide the source of positive feedback. Using a newly developed assay, we find that pBS32 spontaneously cures at a rate of approximately 1 in 3,000 cells stabilized at least in part by the AlfAB active segregation system (17,28-29), and induction of SigN selected for those cells that had spontaneously lost the plasmid. Why and how a plasmid with at least one toxic product on it has been stabilized in the population is unclear, and the selective advantage, if any, of either SigN or its regulon is unknown.

Antisense transcripts that antagonize SigN accumulation also promote stability of the pBS32 plasmid. REND-seq analysis predicted that one or more antisense transcripts read-through *sigN* (25) and here we identify at least two different promoters downstream of, and oriented towards, the *sigN* gene (**Fig 7**). One of the antisense promoters (*P_asiN1_*) appears to overbalance the constitutive sense transcript from *P_sigN2_* and silence *sigN* expression. Thus, production of *sigN* sense transcript must exceed the abundance of the antisense transcript to produce protein. The transition state regulator AbrB represses another equally strong antisense promoter (*P_asiN3_*), either directly or indirectly. AbrB is associated with the repression of a wide variety of genes that become active in post-exponential phase including those producing antimicrobials and biofilm formation (22-24). Here AbrB seems to indirectly promote SigN expression. Why a host cell factor would regulate *sigN* so as to lower the expression threshold and make it easier to initiate a lethal positive feedback loop is unclear. AbrB repression becomes inactivated during the transition to stationary phase (30-32) and this may indicate that whatever the purpose of SigN, it may be selectively antagonized outside of exponential growth.

Many alternative sigma factors, particularly the Sigma 70 ECF subfamily, experience positive feedback and only a few sigmas, such as *B. subtilis* SigM and SigG, have been shown to exhibit deleterious effects on growth when dysregulated (33,34). For deleterious sigmas, it is often to rule out whether one or more targets in the regulon cause toxicity indirectly. Moreover, chromosomally-encoded alternative sigma factors tend to have lower affinity for RNA polymerase core than the vegetative sigma (35-37) and are often co-expressed with anti-sigma factors such that they may be antagonized before, during, and/or after expression (38-40). *Bacillus subtilis* SigN is different in that it is plasmid-encoded, toxic in the absence of its regulon and as yet there is no known anti-sigma factor to hold it in check. Instead, antisense transcripts raise the expression threshold for SigN accumulation, and while perhaps a first for sigma factor control, antisense transcripts are known regulators in horizontally transferred elements such as plasmids, and bacteriophages (41-45). An autoactivating sigma factor with affinity for the RNA core polymerase on par with SigA seems inherently detrimental to the host and it has been speculated that pBS32 is in fact a plasmid prophage like P1 in *E. coli* (5,46). If pBS32 is in fact a prophage in its entirety, override of essential transcription from the host chromosome may be a strategy for phage hyper-proliferation and thus phage survival.

## METHODS

### Strains and growth conditions

*B. subtilis* strains were grown in lysogeny broth (LB) (10 g tryptone, 5 g yeast extract, 5 g NaCl per L) or on LB plates fortified with 1.5% Bacto agar at 37°C. When appropriate, antibiotics were used at the following concentrations: 5 μg/ml kanamycin, 100 μg/ml spectinomycin, 5 μg/ml chloramphenicol, 10 μg/ml tetracycline, and 1 μg/ml erythromycin with 25 μg/ml lincomycin (*mls*). Mitomycin C (MMC, DOT Scientific) was added to the medium at a final concentration when appropriate. Isopropyl β-D-thiogalactopyranoside (IPTG, Sigma) was added to the medium as needed at the indicated concentration.

### Strain Construction

All constructs were first introduced into strains by transformation and then mobilized using SPP1-mediated generalized phage transduction (6,47). All strains used in this study are listed in **Table 2**. All primers used in this study are listed in **Table S1**. All plasmids used are listed in **Table S2**. The abrB::kan allele was obtained from the *Bacillus subtilis* Genetic Stock Center (BGSC, The Ohio State University) (48).

**Table 2:**
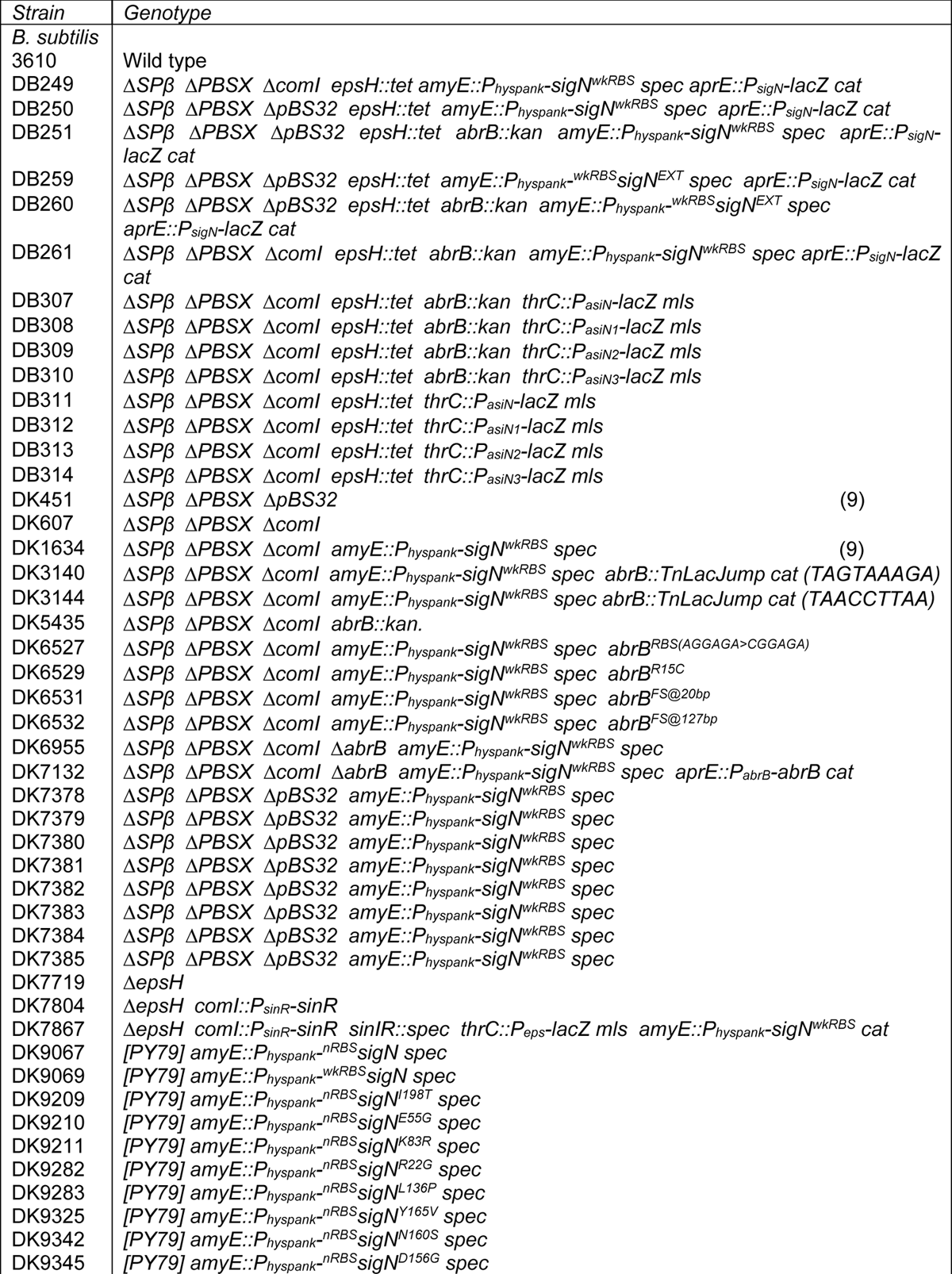
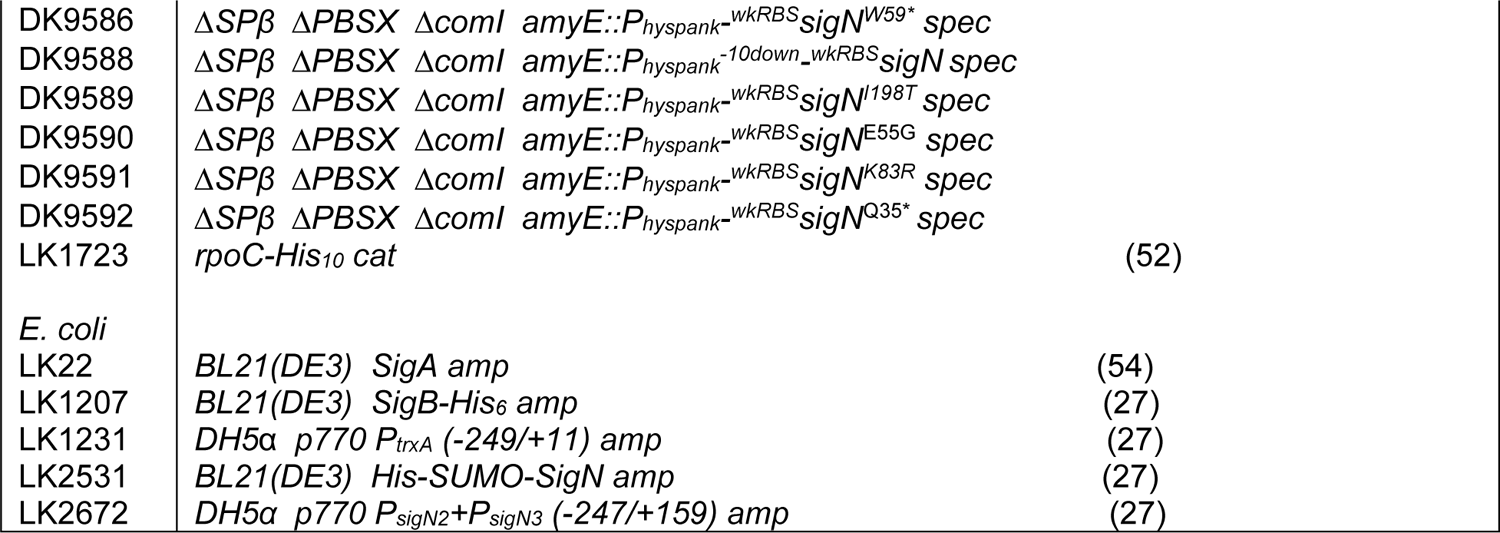
Strains

| Strain | Genotype |
| --- | --- |
| *B. subtilis* |  |
| 3610 | Wild type |
| DB249 | $\Delta SP\beta \Delta PBSX \Delta comI$ *epsH::tet amyE::P<sub>hyspank</sub>-sigN<sup>wkRBS</sup> spec aprE::P<sub>sigN</sub>-lacZ cat* |
| DB250 | $\Delta SP\beta \Delta PBSX \Delta pBS32$ *epsH::tet amyE::P<sub>hyspank</sub>-sigN<sup>wkRBS</sup> spec aprE::P<sub>sigN</sub>-lacZ cat* |
| DB251 | $\Delta SP\beta \Delta PBSX \Delta pBS32$ *epsH::tet abrB::kan amyE::P<sub>hyspank</sub>-sigN<sup>wkRBS</sup> spec aprE::P<sub>sigN</sub>-lacZ cat* |
| DB259 | $\Delta SP\beta \Delta PBSX \Delta pBS32$ *epsH::tet amyE::P<sub>hyspank</sub>-<sup>wkRBS</sup>sigN<sup>EXT</sup> spec aprE::P<sub>sigN</sub>-lacZ cat* |
| DB260 | $\Delta SP\beta \Delta PBSX \Delta pBS32$ *epsH::tet abrB::kan amyE::P<sub>hyspank</sub>-<sup>wkRBS</sup>sigN<sup>EXT</sup> spec aprE::P<sub>sigN</sub>-lacZ cat* |
| DB261 | $\Delta SP\beta \Delta PBSX \Delta comI$ *epsH::tet abrB::kan amyE::P<sub>hyspank</sub>-sigN<sup>wkRBS</sup> spec aprE::P<sub>sigN</sub>-lacZ cat* |
| DB307 | $\Delta SP\beta \Delta PBSX \Delta comI$ *epsH::tet abrB::kan thrC::P<sub>asiN</sub>-lacZ mls* |
| DB308 | $\Delta SP\beta \Delta PBSX \Delta comI$ *epsH::tet abrB::kan thrC::P<sub>asiN1</sub>-lacZ mls* |
| DB309 | $\Delta SP\beta \Delta PBSX \Delta comI$ *epsH::tet abrB::kan thrC::P<sub>asiN2</sub>-lacZ mls* |
| DB310 | $\Delta SP\beta \Delta PBSX \Delta comI$ *epsH::tet abrB::kan thrC::P<sub>asiN3</sub>-lacZ mls* |
| DB311 | $\Delta SP\beta \Delta PBSX \Delta comI$ *epsH::tet thrC::P<sub>asiN</sub>-lacZ mls* |
| DB312 | $\Delta SP\beta \Delta PBSX \Delta comI$ *epsH::tet thrC::P<sub>asiN1</sub>-lacZ mls* |
| DB313 | $\Delta SP\beta \Delta PBSX \Delta comI$ *epsH::tet thrC::P<sub>asiN2</sub>-lacZ mls* |
| DB314 | $\Delta SP\beta \Delta PBSX \Delta comI$ *epsH::tet thrC::P<sub>asiN3</sub>-lacZ mls* |
| DK451 | $\Delta SP\beta \Delta PBSX \Delta pBS32$ (9) |
| DK607 | $\Delta SP\beta \Delta PBSX \Delta comI$ |
| DK1634 | $\Delta SP\beta \Delta PBSX \Delta comI$ *amyE::P<sub>hyspank</sub>-sigN<sup>wkRBS</sup> spec* (9) |
| DK3140 | $\Delta SP\beta \Delta PBSX \Delta comI$ *amyE::P<sub>hyspank</sub>-sigN<sup>wkRBS</sup> spec abrB::TnLacJump cat (TAGTAAAGA)* |
| DK3144 | $\Delta SP\beta \Delta PBSX \Delta comI$ *amyE::P<sub>hyspank</sub>-sigN<sup>wkRBS</sup> spec abrB::TnLacJump cat (TAACCTTAA)* |
| DK5435 | $\Delta SP\beta \Delta PBSX \Delta comI$ *abrB::kan.* |
| DK6527 | $\Delta SP\beta \Delta PBSX \Delta comI$ *amyE::P<sub>hyspank</sub>-sigN<sup>wkRBS</sup> spec abrB<sup>RBS(AGGAGA>CGGAGA)</sup>* |
| DK6529 | $\Delta SP\beta \Delta PBSX \Delta comI$ *amyE::P<sub>hyspank</sub>-sigN<sup>wkRBS</sup> spec abrB<sup>R15C</sup>* |
| DK6531 | $\Delta SP\beta \Delta PBSX \Delta comI$ *amyE::P<sub>hyspank</sub>-sigN<sup>wkRBS</sup> spec abrB<sup>FS@20bp</sup>* |
| DK6532 | $\Delta SP\beta \Delta PBSX \Delta comI$ *amyE::P<sub>hyspank</sub>-sigN<sup>wkRBS</sup> spec abrB<sup>FS@127bp</sup>* |
| DK6955 | $\Delta SP\beta \Delta PBSX \Delta comI \Delta abrB$ *amyE::P<sub>hyspank</sub>-sigN<sup>wkRBS</sup> spec* |
| DK7132 | $\Delta SP\beta \Delta PBSX \Delta comI \Delta abrB$ *amyE::P<sub>hyspank</sub>-sigN<sup>wkRBS</sup> spec aprE::P<sub>abrB</sub>-abrB cat* |
| DK7378 | $\Delta SP\beta \Delta PBSX \Delta pBS32$ *amyE::P<sub>hyspank</sub>-sigN<sup>wkRBS</sup> spec* |
| DK7379 | $\Delta SP\beta \Delta PBSX \Delta pBS32$ *amyE::P<sub>hyspank</sub>-sigN<sup>wkRBS</sup> spec* |
| DK7380 | $\Delta SP\beta \Delta PBSX \Delta pBS32$ *amyE::P<sub>hyspank</sub>-sigN<sup>wkRBS</sup> spec* |
| DK7381 | $\Delta SP\beta \Delta PBSX \Delta pBS32$ *amyE::P<sub>hyspank</sub>-sigN<sup>wkRBS</sup> spec* |
| DK7382 | $\Delta SP\beta \Delta PBSX \Delta pBS32$ *amyE::P<sub>hyspank</sub>-sigN<sup>wkRBS</sup> spec* |
| DK7383 | $\Delta SP\beta \Delta PBSX \Delta pBS32$ *amyE::P<sub>hyspank</sub>-sigN<sup>wkRBS</sup> spec* |
| DK7384 | $\Delta SP\beta \Delta PBSX \Delta pBS32$ *amyE::P<sub>hyspank</sub>-sigN<sup>wkRBS</sup> spec* |
| DK7385 | $\Delta SP\beta \Delta PBSX \Delta pBS32$ *amyE::P<sub>hyspank</sub>-sigN<sup>wkRBS</sup> spec* |
| DK7719 | $\Delta epsH$ |
| DK7804 | $\Delta epsH$ *comI::P<sub>sinR</sub>-sinR* |
| DK7867 | $\Delta epsH$ *comI::P<sub>sinR</sub>-sinR sinIR::spec thrC::P<sub>eps</sub>-lacZ mls amyE::P<sub>hyspank</sub>-sigN<sup>wkRBS</sup> cat* |
| DK9067 | [PY79] *amyE::P<sub>hyspank</sub>-<sup>nRBS</sup>sigN spec* |
| DK9069 | [PY79] *amyE::P<sub>hyspank</sub>-<sup>wkRBS</sup>sigN spec* |
| DK9209 | [PY79] *amyE::P<sub>hyspank</sub>-<sup>nRBS</sup>sigN<sup>I198T</sup> spec* |
| DK9210 | [PY79] *amyE::P<sub>hyspank</sub>-<sup>nRBS</sup>sigN<sup>E55G</sup> spec* |
| DK9211 | [PY79] *amyE::P<sub>hyspank</sub>-<sup>nRBS</sup>sigN<sup>K83R</sup> spec* |
| DK9282 | [PY79] *amyE::P<sub>hyspank</sub>-<sup>nRBS</sup>sigN<sup>R22G</sup> spec* |
| DK9283 | [PY79] *amyE::P<sub>hyspank</sub>-<sup>nRBS</sup>sigN<sup>L136P</sup> spec* |
| DK9325 | [PY79] *amyE::P<sub>hyspank</sub>-<sup>nRBS</sup>sigN<sup>Y165V</sup> spec* |
| DK9342 | [PY79] *amyE::P<sub>hyspank</sub>-<sup>nRBS</sup>sigN<sup>N160S</sup> spec* |
| DK9345 | [PY79] *amyE::P<sub>hyspank</sub>-<sup>nRBS</sup>sigN<sup>D156G</sup> spec* |

|  |  |  |
| --- | --- | --- |
| DK9586 | $\Delta SP\beta$ $\Delta PBSX$ $\Delta comI$ *amyE::P<sub>hyspank</sub><sup>-wkRBS</sup>sigN<sup>W59\*</sup>* *spec* | |
| DK9588 | $\Delta SP\beta$ $\Delta PBSX$ $\Delta comI$ *amyE::P<sub>hyspank</sub><sup>-10down_wkRBS</sup>sigN* *spec* | |
| DK9589 | $\Delta SP\beta$ $\Delta PBSX$ $\Delta comI$ *amyE::P<sub>hyspank</sub><sup>-wkRBS</sup>sigN<sup>I198T</sup>* *spec* | |
| DK9590 | $\Delta SP\beta$ $\Delta PBSX$ $\Delta comI$ *amyE::P<sub>hyspank</sub><sup>-wkRBS</sup>sigN<sup>E55G</sup>* *spec* | |
| DK9591 | $\Delta SP\beta$ $\Delta PBSX$ $\Delta comI$ *amyE::P<sub>hyspank</sub><sup>-wkRBS</sup>sigN<sup>K83R</sup>* *spec* | |
| DK9592 | $\Delta SP\beta$ $\Delta PBSX$ $\Delta comI$ *amyE::P<sub>hyspank</sub><sup>-wkRBS</sup>sigN<sup>Q35\*</sup>* *spec* | |
| LK1723 | *rpoC-His<sub>10</sub>* *cat* | (52) |
| *E. coli* |  |  |
| LK22 | *BL21(DE3)* *SigA* *amp* | (54) |
| LK1207 | *BL21(DE3)* *SigB-His<sub>6</sub>* *amp* | (27) |
| LK1231 | *DH5α* *p770 P<sub>trxA</sub>* (-249/+11) *amp* | (27) |
| LK2531 | *BL21(DE3)* *His-SUMO-SigN* *amp* | (27) |
| LK2672 | *DH5α* *p770 P<sub>sigN2</sub>+P<sub>sigN3</sub>* (-247/+159) *amp* | (27) |

### *LacZ* Reporter Constructs

To generate the β-galactosidase (*lacZ*) reporter *thrC::P_eps_-lacZ mls amp*, the promoter region of *epsA* was amplified with PCR using the primer set 709/3025 from *B. subtilis* 3610 chromosomal DNA. This DNA fragment was digested and cloned with BamHI and EcoRI into pDG1663 (49), which carries an erythromycin-resistance marker and a polylinker upstream of the *lacZ* gene between the two arms of the *thrC* gene to create pDP146.

To generate the pATB28 *thrC::P_asiN_-lacZ mls* reporter construct, the region downstream of *sigN* was PCR amplified from *B. subtilis* 3610 genomic DNA using primers 6576/6577, digested and cloned with BamHI and EcoRI into pDG1663. To generate the pATB51 *thrC::P_asiN1_-lacZ mls* reporter construct, the region downstream of *sigN* was PCR amplified from *B. subtilis* 3610 genomic DNA using primers 6576/6909, digested and cloned with BamHI and EcoRI into pDG1663. To generate the pATB52 *thrC::P_asiN2_-lacZ mls* reporter construct, the region downstream of *sigN* was PCR amplified from *B. subtilis* 3610 genomic DNA using primers 6910/6911, digested and cloned with BamHI and EcoRI into pDG1663. To generate the pATB53 *thrC::P_asiN3_-lacZ mls* reporter construct, the region downstream of *sigN* was PCR amplified from *B. subtilis* 3610 genomic DNA using primers 6912/6577, digested and cloned with BamHI and EcoRI into pDG1663.

### Inducible *^wkRBS^sigN^EXT^* construct

To generate the inducible sigN construct pATB70, the sigN gene plus downstream region were PCR amplified using primer pair 3959/7068 and 3610 genomic DNA as a template. The PCR product was purified, digested with NheI/SphI and cloned into the NheI/SphI sites of plasmid pDR111 containing a polylinker downstream of the Physpank promoter with a spectinomycin resistance cassette all between the arms of the amyE gene (generous gift from David Rudner, Harvard Medical School).

#### Plasmid curing reporter system

To generate the β-galactosidase (*lacZ*) blue-white plasmid curing reporter strain, an Δ*epsH* in-frame markerless deletion construct was generated by transforming pSG37 into DK1042 and selecting for resistance to *mls* at 37° (50). The resulting strain was incubated in 3 ml LB at the permissive temperature for plasmid replication (22°C) for 14h and the culture was serially diluted and plated on LB agar at 37°C. Individual colonies were replica patched onto LB plates and plates containing *mls* to identify *mls*-sensitive colonies that had potentially evicted the plasmid. Colonies that were *mls*-sensitive were screened for smooth colony morphology indicative of a defect in extracellular polysaccharide production and retention of the Δ*epsH* allele (DK7719) was confirmed by PCR length polymorphism using primers 2114/2117.

Next, a markerless *P_sinR_-sinR* complementation construct was introduced into the Δ*epsH* strain DK7719 at the ectopic site *comI* on the pBS32 plasmid. To achieve this, the *comI* 5’ region was PCR amplified with primers 7122/7123 and digested with EcoRI and NheI, and the *comI* 3’ region was amplified with primers 7124/7125 and digested with NheI and SalI. The two fragments were simultaneously ligated into the EcoRI and SalI sites of pMiniMAD (51) to generate pDP532. Next, the *P_sinR_-sinR* region was PCR amplified with primers 7126/7127, digested with NheI and KpnI, and ligated into the NheI and KpnI of the *comI* plasmid pDP532 to create pDP533. Plasmid pDP533 was transformed into DK7719 by transformation and selection for *mls* resistance at 30°C. One colony was re-struck on LB with *mls* at 37°C overnight to force integration of pDP533 into pBS32 and then regrown overnight in 3 ml LB at 30°C to promote pDP533 eviction. The culture was dilution plated for single colonies on LB at 37°C overnight. Retention of *P_sinR_-sinR* at *comI* was confirmed by colony PCR using primers 7122/7125 to generate the markerless *P_sinR_-sinR* integrant DK7804.

Finally, the native copy of *sinIR* was interrupted with a spectinomycin antibiotic cassette (13), a *thrC::P_eps_-lacZ* reporter with an erythromycin resistance cassette, and an *amyE::P_hyspank_-^wkRBS^sigN* construct with a chloramphenicol resistance cassette were sequentially introduced by SPP1-mediated phage transduction to generate DK7867.

### SPP1 Phage Transduction

To a 0.2 ml dense culture grown in TY broth (LB supplemented with 10 mM MgSO_4_ and 100 μM MnSO_4_ after autoclaving), serial dilutions of SPP1 phage stock were added. This mixture was allowed to statically incubate at 37°C for 15 minutes. A 3 ml volume of TYSA (molten TY with 0.5% agar) was added to each mixture and poured on top of fresh TY plates. The plates were incubated at 37 °C overnight. Plates on which plaques formed had the top agar harvested by scraping into a 50 ml conical tube. To release the phage, the tube was vortexed for 20 seconds and centrifuged at 5,000 x *g* for 10 minutes. The supernatant was passed through a 0.45 μm syringe filter and stored at 4 °C.

Recipient cells were grown in 2 ml of TY broth at 37 °C until stationary phase was reached. A 5 µl volume of SPP1 donor phage stock was added to 0.9 of cells and 9 ml of TY broth was added to this mixture. The transduction mixture was allowed to stand statically at room temperature for 30 minutes. After incubation, the mixture was centrifuged at 5,000 x *g* for 10 minutes, the supernatant was discarded, and the pellet was resuspended in the volume left. 100 – 200 µl of the cell suspension was plated on TY fortified with 1.5% agar, 10 mM sodium citrate, and the appropriate antibiotic for selection.

### PCR Amplification

Genomic DNA was isolated from *B. subtilis* strains NCIB 3610, DK6541, DK7383, and DK6527. To amplify the plasmid loci and chromosomal locus the following primer pairs were used: *fliG* (3879/3880), *zpaB* (4762/4763), *zpaQ* (3873/3874), *zpbH* (5566/5567), *zpcJ* (4709/4710), and *zpdN* (5232/5233). Samples were combined according to template and run on a 1% agarose gel for 30 minutes at 100 V.

### Western blotting

*B. subtilis* strains were grown in LB and treated with IPTG or MMC (final concentration 0.3 μg/ml) as reported in (9). Cells were harvested by centrifugation at 1 hour after treatment unless specified. Cells were resuspended to 10 OD_600_ in Lysis buffer [20 mM Tris-HCL (pH 7.0), 10 mM EDTA, 1 mg/ml lysozyme, 10 μg/ml DNAse I, 100 μg/ml RNAse I, 1 mM PMSF] and incubated for 1 hour at 37°C. 20 μl of lysate was mixed with 4 μl 6x SDS loading dye. Samples were separated by 12% sodium dodecyl sulfate-polyacrylamide gel electrophoresis (SDS-PAGE). The proteins were electroblotted onto nitrocellulose and developed with a primary antibody used at a 1:5,000 dilution of anti-SigN, 1:80,000 dilution of anti-SigA, and a 1:10,000 dilution secondary antibody (horseradish peroxidase-conjugated goat anti-rabbit immunoglobulin G). Immunoblot was developed using the Immun-Star HRP developer kit (Bio-Rad).

### β-galactosidase Assay

Biological replicates of *B. subtilis* strains were grown in LB and induced with IPTG to a final concentration of 1mM. Cells grew to an OD_600_ 0.6 and 1 ml was harvested. Cells that contained the plasmid, pBS32, were induced with IPTG and allowed to grow for 1 hour and then harvested by centrifugation. Cells were resuspended in 1 ml of Z-buffer (40 mM NaH_2_PO_4_, 60 mM Na_2_HPO_4_, 1mM MgSO_4_, 10 mM KCl, and 38 mM β-mercaptoethanol) with 0.2 mg/ml of lysozyme and incubated at 30 °C for 15 minutes. Each sample was diluted accordingly with Z-buffer to 500 μl. The reaction was started with 100 μl of 4 mg/ml O-nitrophenyl β-D-galactopyranoside (in Z buffer) and stopped with 1M Na_2_CO_3_ (250 μl). The OD_420_ of each reaction was noted and the β-galactosidase specific activity was calculated using this equation: [OD_420_/(time x OD_600_)] x dilution factor x 1000.

### Protein purification

Wild type RNAP, containing a His10x-tagged β’ subunit was purified from LK1723 as described (52). SigA (LK22) was overproduced and purified as described (53). SigB (LK1207) was overproduced and purified as described (19).

Cells containing the plasmid for overproduction of σ^N^ (LK2531) were grown to OD_600_ ∼0.5 when IPTG was added to a final concentration of 0.3 mM. Cells were then allowed to grow for 3 hours at room temperature, harvested, washed, and resuspended in P buffer (300 mM NaCl, 50 mM Na2HPO4, 3 mM β-mercaptoethanol, 5% glycerol). All purification steps were done in P2 buffer (the same composition as P buffer, but pH 9.5). Cells were then disrupted by sonication and the supernatant was mixed with 1 mL Ni-NTA agarose (QIAGEN) and incubated for 1 h at 4 °C with gentle shaking. Ni-NTA agarose with the bound His-SUMO-SigN was loaded on a Poly-Prep® Chromatography Column (Bio-Rad), washed with P2 buffer and subsequently with the P2 buffer with the 30 mM imidazole. The protein was eluted with P2 buffer containing 400 mM imidazole and fractions containing His-SUMO-SigN were pooled together and dialyzed against P2 buffer. The SUMO tag was subsequently removed by SUMO protease (Invitrogen). The cleavage reaction mixture was then mixed with 1 mL Ni-NTA agarose and allowed to bind for 1 h at 4 °C and centrifuged to pellet the resin. Supernatant was removed, the resin was washed once more with P2 buffer with 3 mM β-ME. The supernatants (containing SigN) were pooled together and dialysed against storage P2 buffer (P2 buffer and 50% glycerol). The protein was stored at −20 °C.

### Transcription *in vitro*

Multiple round transcriptions *in vitro* were performed with the *B. subtilis* RNAP core reconstituted with SigA (ratio 1:1) and either without or with increasing amounts of SigN or SigB (ratios 1:0.5, 1:1, 1:2, 1:4, 1:8; RNAP was 1) in storage buffer (50 mM Tris-HCl, pH 8.0, 0.1 M NaCl, 50% glycerol) for 30 minutes at 30°C (27,54). First, the two sigma factors were mixed together and then the RNAP core was added. The obtained holoenzymes reflected the competition between these sigma factors for the core. Reconstituted holoenzymes were then used in multiple round transcription reactions in 10 µl reaction volumes. The transcription buffer contained 40 mM Tris-HCl pH 8.0, 10 mM MgCl2, 1 mM dithiothreitol (DTT), 0.1 mg/mL BSA, 150 mM KCl, and all four NTPs (200 μM ATP, CTP, GTP each) plus 10 μM UTP and 2 µM radiolabeled [α-^32^P] UTP. Every reaction also contained supercoiled DNA template (100 ng/reaction). The LK2672 plasmid containing P*_sigN2_*+P*_sigN3_* was used for competition experiments. As a control to verify that SigB was active, LK1231 (P*_trxA_*) was used. The transcriptions were initiated with 30 nM RNAP holoenzyme (final concentration), allowed to proceed for 15 minutes at 37 °C and stopped with equal volumes of formamide stop solution (95% formamide, 20 mM EDTA, pH 8.0). Samples were loaded onto 7 M urea-7% polyacrylamide gel and electrophoresis was performed. The gels were dried and scanned with Amersham Typhoon (Cytiva), visualization and analysis were made using the Quantity One software (Bio-Rad).

## ACKNOWLEDGEMENTS

Felix Dempwolff, Kate Hummels, Bat-Erdene Magyarjav, Reid Oshiro, and Sundharraman Subramanian for technical support. This work was supported by NIH Grant R35 GM131783 to DBK, Czech Science Foundation Grant 22-06342K to LK and the project European Union - Next Generation EU National Institute of Virology and Bacteriology Programme EXCELES Project LX22NPO5103 to LK.

